# Reproducibility Assessment of Neuromelanin-Sensitive Magnetic Resonance Imaging Protocols for Region-of-Interest and Voxelwise Analyses

**DOI:** 10.1101/781815

**Authors:** Kenneth Wengler, Xiang He, Anissa Abi-Dargham, Guillermo Horga

**Author notes:** These authors contributed equally (Anissa Abi-Dargham, Guillermo Horga). Corresponding Author: Kenneth Wengler, PhD, Department of Psychiatry, Columbia University, Division of Translational Imaging, New York State Psychiatric Institute, 1051 Riverside Dr, Unit 31, New York, NY 10032.

## Abstract

Neuromelanin-sensitive MRI (NM-MRI) provides a noninvasive measure of the content of neuromelanin (NM), a product of dopamine metabolism that accumulates with age in dopamine neurons of the substantia nigra (SN). NM-MRI has been validated as a measure of both dopamine neuron loss, with applications in neurodegenerative disease, and dopamine function, with applications in psychiatric disease. Furthermore, a voxelwise-analysis approach has been validated to resolve substructures, such as the ventral tegmental area (VTA), within midbrain dopaminergic nuclei thought to have distinct anatomical targets and functional roles. NM-MRI is thus a promising tool that could have diverse research and clinical applications to noninvasively interrogate in vivo the dopamine system in neuropsychiatric illness. Although a test-retest reliability study by Langley et al. using the standard NM-MRI protocol recently reported high reliability, a systematic and comprehensive investigation of the performance of the method for various acquisition parameters and preprocessing methods has not been conducted. In particular, most previous studies used relatively thick MRI slices (∼3 mm), compared to the typical in-plane resolution (∼0.5 mm) and to the height of the SN (∼15 mm), to overcome technical limitations such as specific absorption rate and signal-to-noise ratio, at the cost of partial-volume effects. Here, we evaluated the effect of various acquisition and preprocessing parameters on the strength and test-retest reliability of the NM-MRI signal to determine optimized protocols for both region-of-interest (including whole SN/VTA-complex and atlas-defined dopaminergic nuclei) and voxelwise measures. Namely, we determined a combination of parameters that optimizes the strength and reliability of the NM-MRI signal, including acquisition time, slice-thickness, spatial-normalization software, and degree of spatial smoothing. Using a newly developed, detailed acquisition protocol, across two scans separated by 13 days on average, we obtained intra-class correlation values indicating excellent reliability and high contrast-to-noise, which could be achieved with a different set of parameters depending on the measures of interest and experimental constraints such as acquisition time. Based on this, we provide detailed guidelines covering acquisition through analysis and recommendations for performing NM-MRI experiments with high quality and reproducibility. This work provides a foundation for the optimization and standardization of NM-MRI, a promising MRI approach with growing applications throughout clinical and basic neuroscience.

**Highlights:**

- A detailed NM-MRI volume placement protocol is described.
- Guidelines covering acquisition through analysis for NM-MRI are given.
- A test-retest study in 10 healthy subjects shows high reproducibility for region-of-interest (ROI) and voxelwise analyses.
- ∼3 minutes of NM-MRI data is needed for high-quality ROI-analysis.
- ∼6 minutes of NM-MRI data is needed for high-quality voxelwise-analysis.

## 1. Introduction

Neuromelanin (NM) is an insoluble dark pigment that consists of melanin, proteins, lipids, and metal ions (Zecca et al., 2008). Neurons containing NM are present in specific brain regions of the human central nervous system with particularly high concentrations found in the dopaminergic neurons of the substantia nigra (SN) and noradrenergic neurons of the locus coeruleus (LC) (Zecca et al., 1996; Zucca et al., 2014). NM is synthesized by iron-dependent oxidation of dopamine, norepinephrine, and other catecholamines in the cytosol, to semi-quinones and quinones (Sulzer and Zecca, 1999). While initially present in the cytosol, NM accumulates within cytoplasmic organelles via macroautophagy that results in the undegradable material being taken into autophagic vacuoles (Sulzer et al., 2000). These vacuoles then fuse with lysosomes and other autophagic vacuoles containing lipid and protein components to form the final NM-containing organelles (Zucca et al., 2014). These organelles contain NM pigment along with metals, abundant lipid bodies, and protein matrix (Zecca et al., 2000; Zucca et al., 2018). This process was shown to be driven by excess cytosolic catecholamines, such as that resulting from L-DOPA exposure, that are not accumulated in synaptic vesicles and can be inhibited by treatment with the iron chelator desferrioxamine (Cebrián et al., 2014; Sulzer et al., 2000). NM-containing organelles first appear in humans between 2 and 3 years of age (Cowen, 1986) and gradually accumulate with age (Zecca et al., 2008; Zucca et al., 2018).

The paramagnetic nature of the NM-iron complexes within the NM-containing organelles (Zecca et al., 1996; Zecca et al., 2004) enables them to be noninvasively imaged using magnetic resonance imaging (MRI) (Cassidy et al., 2019; Sasaki et al., 2006; Sulzer et al., 2018; Trujillo et al., 2017). NM-sensitive MRI (NM-MRI) produces hyperintense signals in neuromelanin-containing regions such as the SN and LC due to the short longitudinal relaxation time (T_1_) of the NM-complexes and saturation of the surrounding white matter (WM) by either direct magnetization transfer (MT) pulses (Chen et al., 2014) or indirect MT effects (Sasaki et al., 2006) (see Trujillo et al. for a detailed investigation of NM-MRI contrast mechanisms (Trujillo et al., 2017)). While most previous NM-MRI studies have used indirect MT effects, images with direct MT pulses achieve greater sensitivity (Langley et al., 2015; Schwarz et al., 2013) and were recently shown to be directly related to NM concentration (Cassidy et al., 2019). NM-MRI has also been validated as a measure of dopaminergic neuron loss in the SN (Kitao et al., 2013) and several studies have shown that this method can capture the well-known loss of NM-containing neurons in the SN of individuals with Parkinson’s disease (Sulzer et al., 2018). More recently, NM-MRI was validated as a marker of dopamine function, with the NM-MRI signal in the SN demonstrating a significant relationship to Positron emission tomography (PET) measures of dopamine release capacity in the striatum (Cassidy et al., 2019). Furthermore, a voxelwise-analysis approach was validated to resolve substructures within dopaminergic nuclei thought to have distinct anatomical targets and functional roles (Cassidy et al., 2019; Haber et al., 1995; Roeper, 2013; Weinstein et al., 2017). This voxelwise approach may thus allow for a more anatomically precise interrogation of specific midbrain circuits encompassing subregions within the SN or small nuclei such as the ventral tegmental area (VTA), which may in turn increase the accuracy of NM-MRI markers for clinical or mechanistic research. For example, voxelwise NM-MRI may facilitate investigations into the specific subregions within the SN/VTA-complex projecting to the head of the caudate, which are of particular relevance in the study of psychosis (Weinstein et al., 2017), or help capture the known topography of SN neuronal loss in Parkinson’s disease (Cassidy et al., 2019; Damier et al., 1999; Fearnley and Lees, 1991). An additional benefit of the voxelwise-analysis is avoiding the circularity that can incur when defining ROIs based on the NM-MRI images that are then used to read out the signal in those same regions. Most previous studies have used the high signal region in the NM-MRI images to define the SN region-of-interest (ROI) that is used for further analysis. While this may be appropriate if the goal of the study is to measure the volume of the SN, it can be problematic for analysis of the contrast-to-noise ratio (CNR) because the selected ROI is biased towards high CNR voxels.

NM-MRI can be a promising tool with diverse research and clinical applications to noninvasively interrogate in vivo the dopamine system. However, this is critically dependent on a thorough investigation of the performance of the method for various acquisition parameters and preprocessing methods. In particular, most previous studies used relatively thick MRI slices (∼3 mm) (Langley et al., 2017; Sasaki et al., 2008; Schwarz et al., 2011), compared to the in-plane resolution (∼0.5 mm) and to the height of the SN (∼15 mm) (Pauli et al., 2018), to overcome technical limitations such as specific absorption rate and signal-to-noise ratio (SNR), at the cost of partial-volume effects. Additionally, previous studies acquired multiple measurements that were subsequently averaged to improve the SNR of the technique with the expense of increased scanning time. Although time in the MRI is expensive and should be minimized to improve patient compliance, a detailed investigation into how many measurements are necessary for robust NM-MRI has not been reported. A recent study reported high reproducibility for ROI-analysis (Langley et al., 2017). However, this study did not provide a detailed description of NM-MRI volume placement and the two scans were obtained within a single session, although subjects were removed and repositioned in-between scans.

Here, we evaluate the effect of various acquisition and preprocessing parameters on the strength and test-retest reliability of the NM-MRI signal to determine optimized protocols for both ROI and voxelwise measures. Three new NM-MRI sequences with slice-thickness of 1.5 mm, 2 mm, and 3 mm were compared to the literature standard sequence with 3 mm slice-thickness (Cassidy et al., 2019; Chen et al., 2014). Using our step-by-step acquisition protocol, across the two acquired scans we obtained intra-class correlation coefficient (ICC) values indicating excellent reliability and high CNR, which could be achieved with a different set of parameters depending on the measures of interest and experimental constraints such as acquisition time. A detailed analysis of the CNR and ICC provide evidence for the optimal spatial-normalization software, number of measurements (acquisition time), slice thickness, and spatial smoothing. Based on this, we provide detailed guidelines covering acquisition through analysis and recommendations for performing NM-MRI experiments with high quality and reproducibility.

## 2. Methods

### 2.1. Participants

10 healthy subjects underwent 2 MRI exams (test and re-test) on a 3T Prisma MRI (Siemens, Erlangen, Germany) using a 64-channel head coil. The test-retest scans were separated by a minimum of 2 days. Inclusion criteria were: age between 18 and 65 years and no MRI contraindications. Exclusion criteria were: history of neurological or psychiatric diseases, pregnancy or nursing, and inability to provide written consent. This study was approved by the Stony Brook University Institutional Review Board and written informed consent was obtained from all subjects.

### 2.2. Magnetic resonance imaging

A T1-weighted (T1w) image was acquired for processing of the NM-MRI image using a 3D magnetization prepared rapid acquisition gradient echo (MPRAGE) sequence with the following parameters: spatial resolution = 0.8 × 0.8 × 0.8 mm^3^; field-of-view (FOV) = 166 × 240 × 256 mm^3^; echo time (TE) = 2.24 ms; repetition time (TR) = 2,400 ms; inversion time (TI) = 1060 ms; flip angle = 8°; in-plane acceleration, GRAPPA = 2 (Griswold et al., 2002); bandwidth = 210 Hz/pixel. A T2-weighted (T2w) image was acquired for processing of the NM-MRI image using a 3D sampling perfection with application-optimized contrasts by using flip angle evolution (SPACE) sequence with the following parameters: spatial resolution = 0.8 × 0.8 × 0.8 mm^3^; FOV = 166 × 240 × 256 mm^3^; TE = 564 ms; TR = 3,200 ms; echo spacing = 3.86 ms; echo train duration = 1,166 ms; variable flip angle (T2 var mode); in-plane acceleration = 2; bandwidth = 744 Hz/pixel. NM-MRI images were acquired using 4 different gradient 2D recalled echo sequences with magnetization transfer contrast (2D GRE-MTC) (Chen et al., 2014). The following parameters were consistent across the 4 2D GRE-MT sequences: in-plane resolution = 0.4 × 0.4 mm^2^; FOV = 165 × 220 mm^2^; flip angle = 40°; slice gap = 0 mm; bandwidth = 390 Hz/pixel; MT frequency offset = 1.2 kHz; MT pulse duration = 10 ms; MT flip angle = 300°; partial k-space coverage of MT pulse as described by Chen et al. (Chen et al., 2014). The partial k-space coverage MT pulses were applied in a trapezoidal fashion as described by Parker et al. (Parker et al., 1995) with ramp-up and ramp-down coverage of 20% and plateau coverage of 40%. Other 2D GRE-MTC sequence parameters that differed across the 4 sequences are listed in Table 1. The order of the 4 NM-MRI sequences was randomized across all subjects and sessions.

**Table 1.** 2D GRE-MTC Sequence Parameters Used for NM-MRI.

| Sequence | TE (ms) | TR (ms) | Slice-Thickness (mm) | Number of Slices | Number of Acquisitions | Acquisition Time (minutes:s) | In-Plane Acceleration |
| --- | --- | --- | --- | --- | --- | --- | --- |
| NM-1.5 mm | 4.11 | 444 | 1.5 | 16 | 11 | 10:51 | 2 |
| NM-2 mm | 3.61 | 321 | 2 | 12 | 15 | 10:42 | 2 |
| NM-3 mm | 3.61 | 214 | 3 | 8 | 23 | 10:56 | 2 |
| NM-3 mm Standard | 3.87 | 273 | 3 | 10 | 10 | 11:02 | None |

### 2.3. Neuromelanin-MRI placement protocol

Given the limited coverage of the NM-MRI protocol in the inferior-superior direction (∼30 mm), a detailed NM-MRI volume placement procedure based on distinct anatomical landmarks was developed to improve within-subject and across-subject repeatability. The placement protocol makes use of the sagittal, coronal, and axial 3D T1w images. Furthermore, the coronal and axial images were reformatted along the anterior commissure-posterior commissure (AC-PC) line. The following is the step-by-step procedure used for NM-MRI volume placement:

1. Identification of the sagittal image showing the greatest separation between the midbrain and thalamus (Fig. 1A).
2. Using the sagittal image from the end of Step 1, finding the coronal plane that identifies the most anterior aspect of the midbrain (Fig. 1B).
3. Using the coronal image from the end of Step 2, finding the axial plane that identifies the inferior aspect of the third ventricle (Fig. 1C and Fig. 1D).
4. Setting the superior boundary of the NM-MRI volume to be 3 mm superior to the axial plane from the end of Step 3 (Fig. 1E).

An example of the final placement from a representative subject is shown in Fig. 2.

**Fig. 1:**
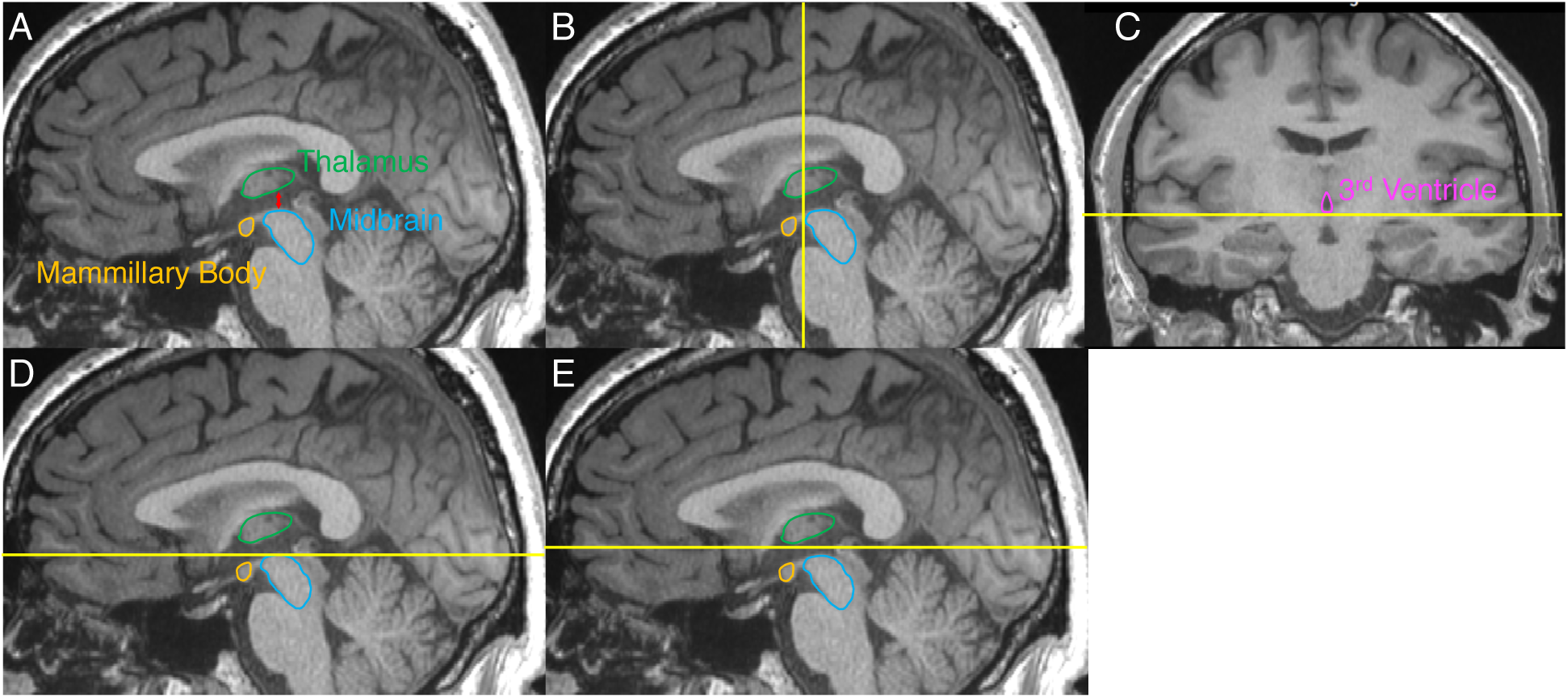
Illustration of the step-by-step procedure for placement of the NM-MRI volume. Yellow lines indicate the position of slices used for placing the volume. (A) Sagittal image showing the greatest separation between the midbrain and thalamus. (B) Coronal plane that identifies the most anterior aspect of the midbrain on the image from A. (C) Axial plane that identifies the inferior aspect of the third ventricle on the image of the coronal plane from B. (D) Location of the axial plane from C identified on the image from A. (E) The axial plane denoting the superior boundary of the NM-MRI volume. This axial plane is the axial plane from D shifted superiorly 3 mm.

**Fig. 2:**
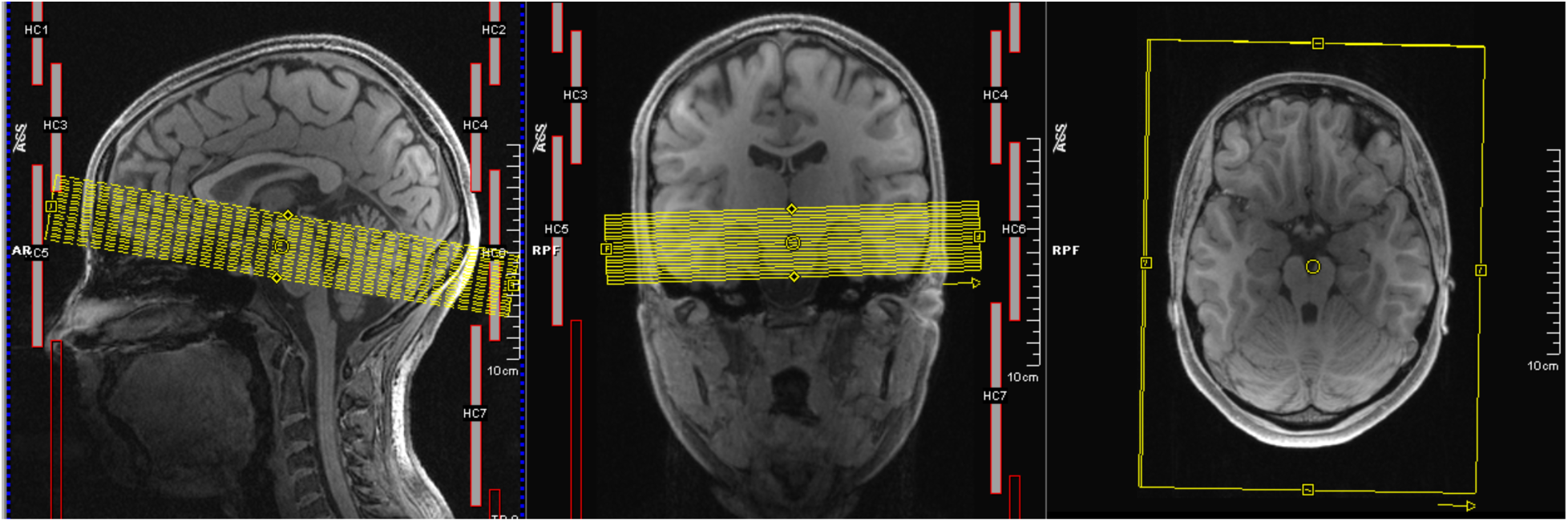
Final NM-MRI volume placement from a representative subject.

### 2.4. Neuromelanin-MRI preprocessing

The NM-MRI preprocessing pipeline was written in Matlab (MathWorks, Natick, MA) using SPM12 (Penny et al., 2011). The steps of the pipeline were as follows: 1) The intra-sequence acquisitions were realigned to the first acquisition to correct for inter-acquisition motion; 2) The motion corrected NM-MRI images were subsequently averaged; 3) The average NM-MRI images were then coregistered to the T1w image; 4) The T1w image was spatially normalized to a standard MNI template using 4 different software: ANTs (Avants et al., 2008; Avants et al., 2009), FSL (Andersson et al., 2007; Jenkinson et al., 2012), SPM12’s Unified Segmentation (referred to as SPM12 throughout) (Ashburner and Friston, 2005; Penny et al., 2011) and SPM12’s DARTEL (referred to as DARTEL throughout) (Ashburner, 2007; Penny et al., 2011). The warping parameters to normalize the T1w image to the MNI template were then applied to the coregistered NM-MRI images using the respective software. The resampled resolution of the spatially normalized NM-MRI images was 1 mm, isotropic. Additionally, the spatially normalized NM-MRI images were spatially smoothed using 3D Gaussian kernels with full-width-at-half-maximum (FWHM) of 0 mm (no smoothing), 1 mm, 2 mm, and 3 mm. All analyses using manually traced ROIs used the standard 1 mm spatial smoothing. All ROI-analysis results used 0 mm of spatial smoothing and all voxelwise-analysis results used 1 mm of spatial smoothing unless otherwise specified.

### 2.5. Neuromelanin-MRI analysis

The NM-MRI contrast-to-noise ratio (CNR) at each voxel *V* was calculated as the relative change in NM-MRI signal intensity *I* from a reference region *RR* of white matter tracts known to have minimal NM content, the crus cerebri (CC) (Cassidy et al., 2019), as *CNR*_*V*_ *=* [*I* – *mode*(*I*_*RR*_)]/*mode(I*_*RR*_).

Two-way mixed, single score intraclass correlation coefficient [ICC(3,1)] (Shrout and Fleiss, 1979) were used to assess the test-retest reliability of NM-MRI. This ICC is a measure of consistency between the first and second measurements that does not penalize consistent changes across all subjects (e.g., if the retest CNR is consistently higher than the test CNR for all subjects). The maximum ICC is 1, indicating perfect reliability, ICC over 0.75 indicates “excellent” reliability, ICC between 0.75 and 0.6 indicates “good” reliability, ICC between 0.6 and 0.4 indicates “fair” reliability, and ICC under 0.4 indicates “poor” reliability (Cicchetti, 1994). ICC(3,1) values were calculated for three conditions: the average CNR within a given ROI (1 ICC value per ROI; ICC_ROI_); the across-subject voxelwise CNR (1 ICC value per voxel; ICC_ASV_); the within-subject voxelwise CNR (1 ICC value per subject; ICC_WSV_). ICC_ROI_ provides a measure of the reliability of the average CNR within an ROI across all subjects, thus providing a measure of the reliability of the ROI-analysis approach. ICC_ASV_ provides a measure of the reliability of CNR_V_ at each voxel within an ROI across all subjects, thus providing a measure of the reliability of the voxelwise-analysis approach. ICC_WSV_ provides a measure of the reliability of the spatial pattern of CNR_V_ across voxels within each of the subjects individually, which provides a complementary measure of reliability of the voxelwise-analysis approach.

The ROIs used in this study included a manually traced mask of the SN defined from a previous study (Cassidy et al., 2019) and ROIs of the SN/VTA-complex nuclei: SN pars compacta (SNc), SN pars reticulata (SNr), ventral tegmental area (VTA), and parabrachial pigmented nucleus (PBP) as defined from a high-resolution probabilistic atlas (Pauli et al., 2018). Fig 3 shows the ROIs overlaid on a template NM image.

**Fig. 3:**
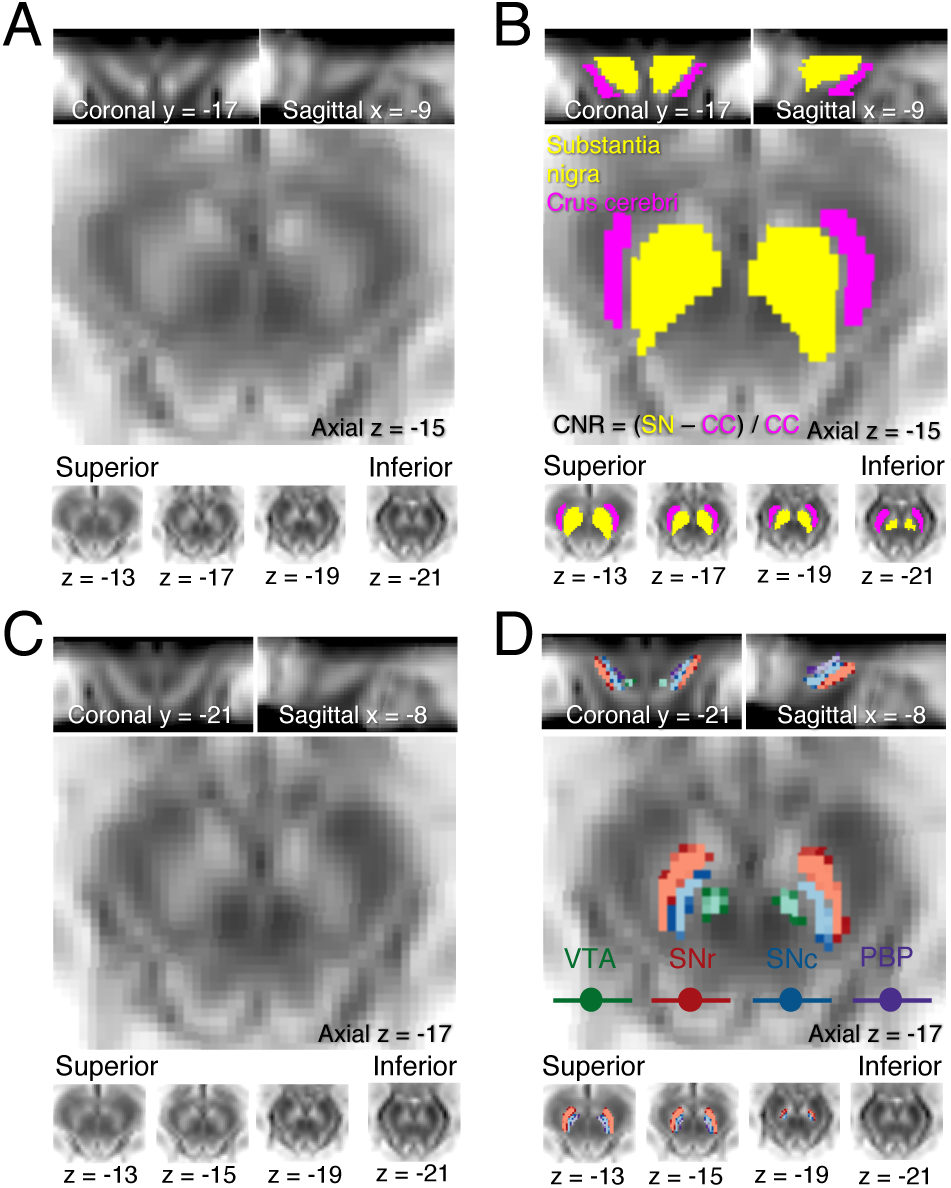
Spatial normalization and anatomical masks for analysis of NM-MRI images. (A) Average NM-MRI image created by averaging the spatially normalized NM-MRI images from 10 individuals in MNI space. Note the high signal intensity in the SN. (B) Masks for the SN (yellow voxels) and the CC (pink voxels) reference region (used in the calculation of CNR) are overlaid onto the template in A. These anatomical masks were made by manual tracing on a NM-MRI template from a previous study (Cassidy et al., 2019). (C) The same average NM-MRI image from A. (D) Probabilistic masks for the VTA, SNr, SNc, and PBP as defined from a high-resolution probabilistic atlas (Pauli et al., 2018) overlaid onto the template in C. The color scaling for probabilistic masks goes from P = 0.5 (darkest) to P = 0.8 (lightest).

## 3. Results

### 3.1. Demographics

The test-retest MRI exams were separated by 13 ± 13 (mean ± standard deviation) days on average with a median of 8 days, minimum of 2 days, and maximum of 38 days. Of the 10 subjects, 4 were male and 6 were female; 4 were Caucasian and 6 were Asian; 9 were right-handed and 1 was left-handed. The average age was 27 ± 5 years (mean ± standard deviation). None of the subjects reported current cigarette smoking or recreational drug use.

### 3.2. Required acquisition time

Plots of ICC_ROI_ and CNR_ROI_ within the manually traced mask as a function of acquisition time for each of the NM-MRI sequences and spatial normalization software are shown in Fig. 4. In general, all NM-MRI sequences and spatial normalization software achieved excellent test-retest reliability within 3 minutes of acquisition time and CNR_ROI_ was not affected by acquisition time. The NM-1.5 mm sequence had the highest CNR_ROI_ for all spatial normalization software while the NM-3 mm sequence had the lowest.

**Fig. 4:**
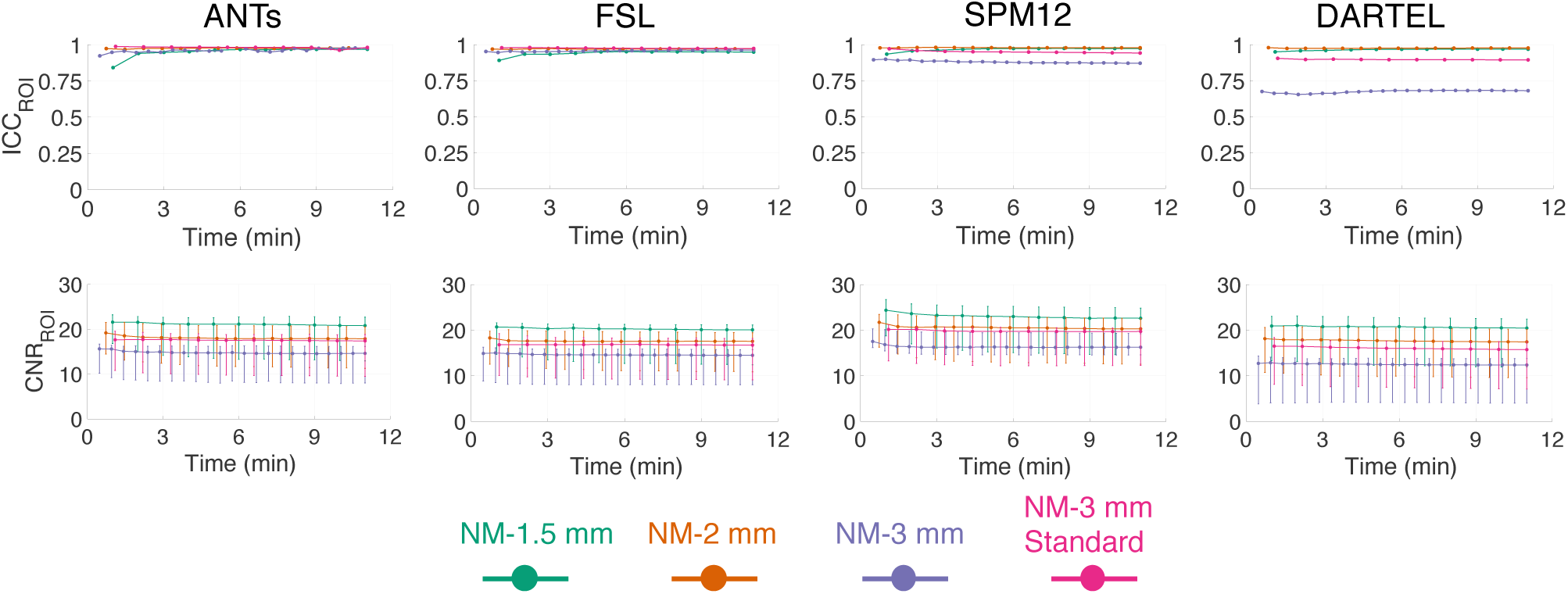
ICC_ROI_ (top row) and CNR_ROI_ (bottom row) within the manually traced mask of the SN/VTA-complex (Fig. 3B) as a function of acquisition time. Data points denote the median and error bars indicate the 25^th^ and 75^th^ percentiles.

Plots of the ICC_ASV_, ICC_WSV_, and CNR_V_ within the manually traced mask as a function of acquisition time for each of the NM-MRI sequences and spatial normalization software are shown in Fig. 5. In general, all spatial normalization software and NM-MRI sequences except for NM 3-mm achieved excellent test-retest reliability within 6 minutes of acquisition time and CNR_ROI_ was not affected by acquisition time. The NM-1.5 mm sequence had the highest CNR_ROI_ for all spatial normalization software while the NM-3 mm sequence had the lowest.

**Fig. 5:**
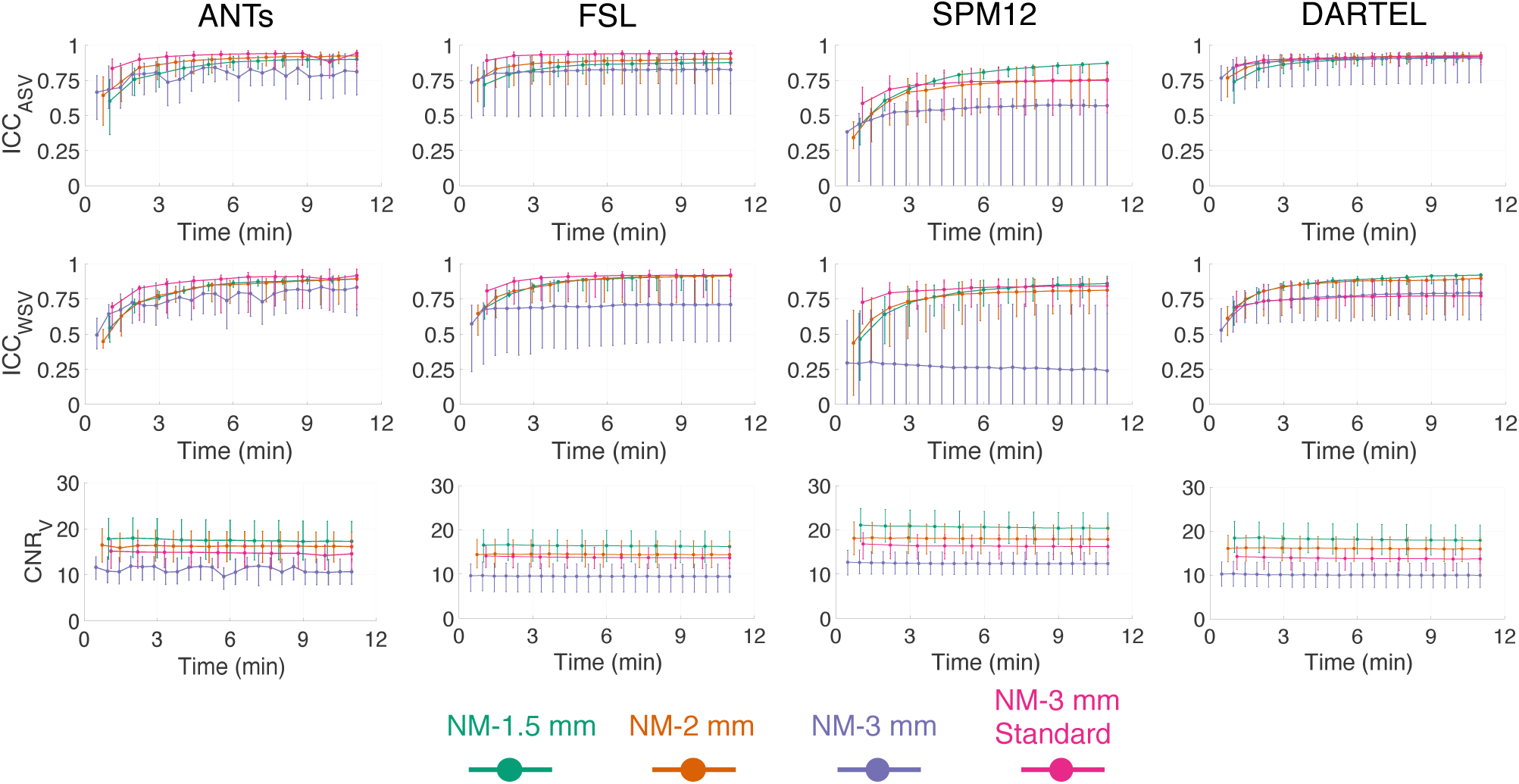
ICC_ASV_ (top row), ICC_WSV_ (middle row), and CNR_V_ (bottom row) of voxels within the manually traced mask of the SN/VTA-complex (Fig. 3B) as a function of acquisition time. Data points denote the median and error bars indicate the 25^th^ and 75^th^ percentiles.

### 3.3. Choice of NM-MRI sequence

Scatterplots of the ICC_ASV_ and CNR_V_ of each voxel within the manually traced mask for each of the NM-MRI sequences and spatial normalization software are shown in Fig. 6. Table 2 lists the 25^th^ percentile, median, and 75^th^ percentile of ICC_ASV_ and CNR_V_ values shown in Fig 6. The NM-1.5 mm sequence consistently showed the highest CNR_V_, greatest spread in CNR_V_, lowest correlation between CNR_V_ and ICC_ASV_, and high ICC_ASV_ across all spatial normalization software. Because the NM-1.5 mm sequence demonstrated the best performance, further optimization was performed for this sequence and the following sections only use data from this sequence.

**Table 2:** ICC_ASV_, CNR_V_, and Spearman’s rho of their relationship for each NM-MRI sequence and spatial normalization software. ICC_ASV_ and CNR_V_ values are from within the manually traced mask of the SN/VTA-complex (Fig. 3B) and are listed as 25^th^ percentile, median, 75^th^ percentile. Spearman’s rho values represent the relationship between ICC_ASV_ and CNR_V_ of voxels within the manually traced mask.

| Sequence | Spatial Normalization Software | ICC<sub>ASV</sub> | CNR<sub>V</sub> | $\rho$ |
| --- | --- | --- | --- | --- |
| NM-1.5 mm | ANTs | 0.86, 0.91, 0.94 | 13.2, 17.3, 21.6 | 0.26 |
|  | FSL | 0.82, 0.88, 0.92 | 12.6, 16.2, 19.7 | 0.22 |
|  | SPM12 | 0.77, 0.86, 0.91 | 16.7, 20.4, 23.8 | 0.26 |
|  | DARTEL | 0.85, 0.89, 0.92 | 15.3, 18.8, 22.3 | 0.25 |
| NM-2 mm | ANTs | 0.87, 0.92, 0.95 | 12.7, 16.1, 19.3 | 0.37 |
|  | FSL | 0.73, 0.90, 0.95 | 11.5, 14.4, 17.5 | 0.49 |
|  | SPM12 | 0.65, 0.81, 0.90 | 14.5, 17.9, 21.1 | 0.35 |
|  | DARTEL | 0.81, 0.88, 0.92 | 14.3, 17.3, 19.9 | 0.26 |
| NM-3 mm | ANTs | 0.21, 0.77, 0.93 | 7.9, 10.6, 13.4 | 0.42 |
|  | FSL | 0.51, 0.83, 0.94 | 5.9, 9.5, 12.2 | 0.33 |
|  | SPM12 | -0.04, 0.24, 0.91 | 9.9, 12.4, 14.9 | 0.27 |
|  | DARTEL | 0.32, 0.80, 0.86 | 7.3, 10.2, 12.9 | 0.54 |
| NM-3 mm Standard | ANTs | 0.84, 0.95, 0.97 | 11.3, 14.6, 17.1 | 0.30 |
|  | FSL | 0.86, 0.94, 0.96 | 11.3, 13.7, 15.9 | 0.35 |
|  | SPM12 | 0.70, 0.84, 0.89 | 13.2, 16.2, 18.6 | 0.42 |
|  | DARTEL | 0.85, 0.91, 0.94 | 10.5, 13.2, 15.3 | 0.47 |

**Fig. 6:**
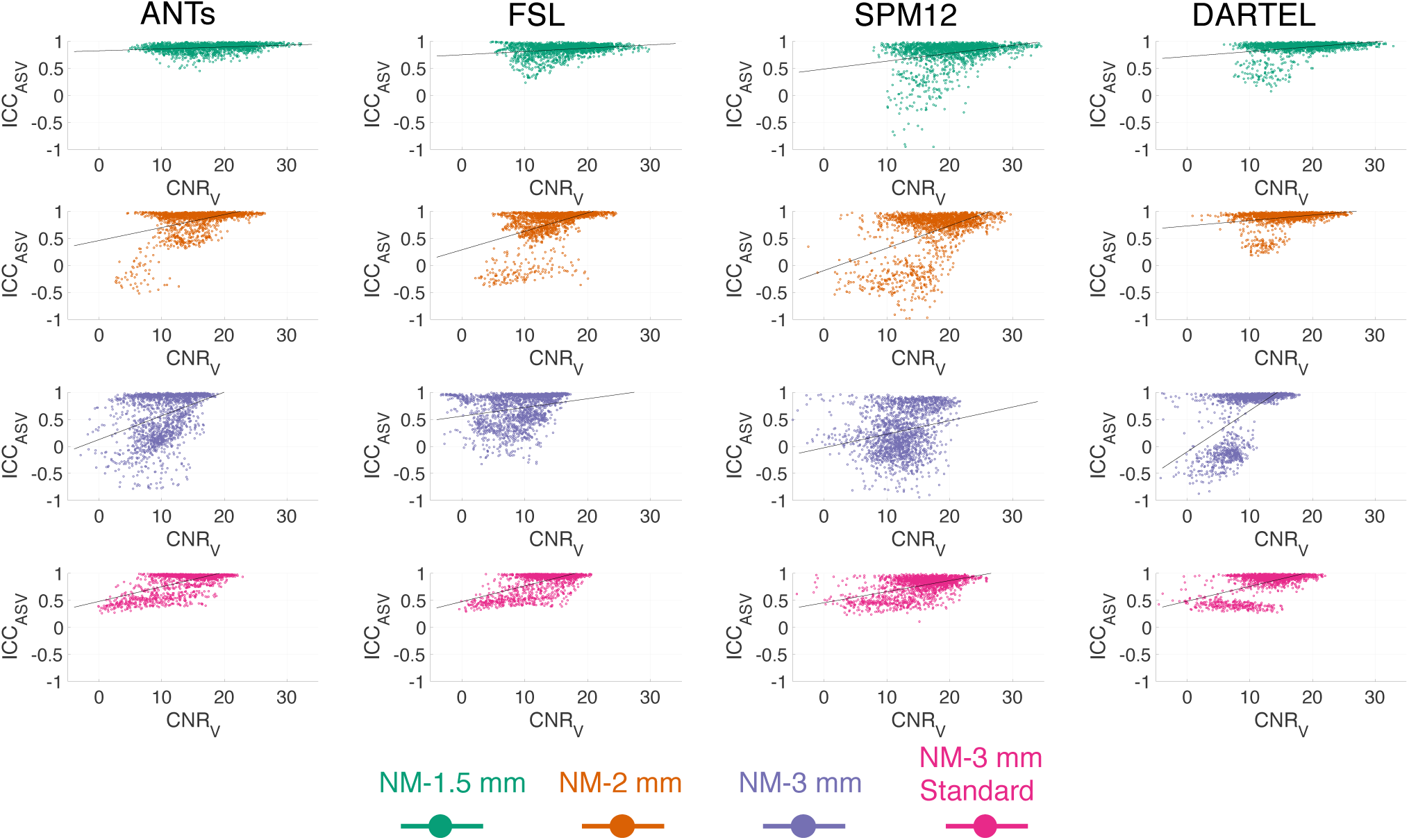
Scatterplots of ICC_ASV_ and CNR_V_ for each of the NM-MRI sequences and spatial normalization software. Each data point represents one voxel within the manually traced mask of the SN/VTA-complex (Fig. 3B). The solid lines indicate the linear fit of the relationship between ICC_ASV_ and CNR_V_.

### 3.4. Choice of spatial normalization software

To determine which spatial normalization software should be used for voxelwise-analysis of NM-MRI data, multiple linear regression analysis predicting ICC_ASV_ of voxels within the manually traced mask as a function of their coordinates in *x* (absolute distance from the midline), *y*, and *z* directions was used. The rationale here is that, with an optimal method, the ICC should be highest and homogeneous across voxels such that the voxel’s anatomical location should not predict its associated ICC value. This analysis showed that ANTs achieved the best performance in that anatomical position was least predictive of ICC_ASV_ and it provided the highest ICC_ASV_ (Fig. 7). This result was consistent with a previous study where ANTs outperformed 13 other spatial-normalization algorithms (Klein et al., 2009).

**Fig. 7:**
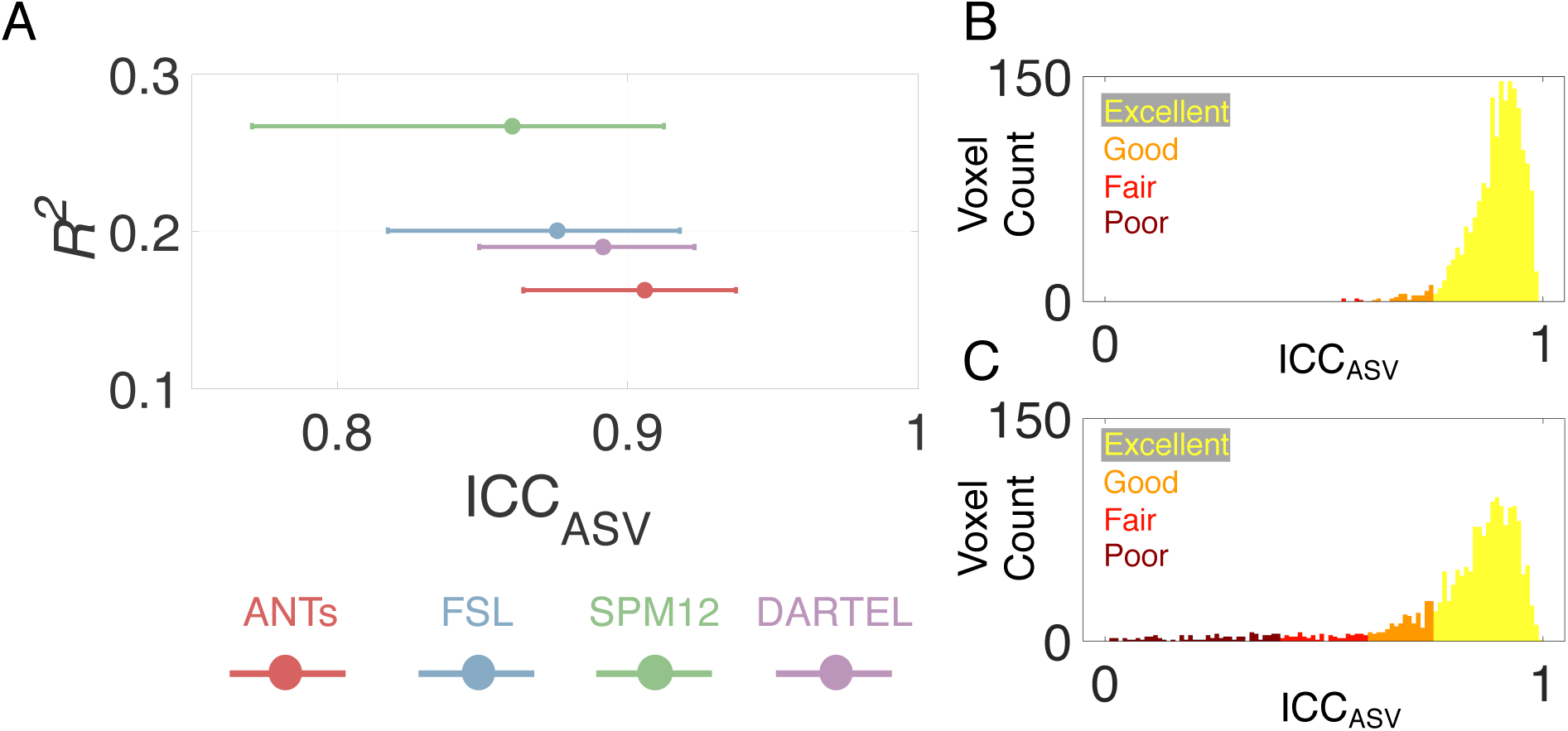
(A) Predictive value (*R*^*2*^) of anatomical position on ICC_ASV_ and ICC_ASV_ of voxels within the manually traced mask of the SN/VTA-complex (Fig. 3B) for NM-1.5 mm sequence and each of the spatial normalization software. Data points denote the median and error bars indicate the 25^th^ and 75^th^ percentiles. (B) Histogram of ICC_ASV_ of voxels within the manually traced mask for NM-1.5 mm sequence and ANTs spatial normalization software, which is the best performing method as per A. (C) Histogram of ICC_ASV_ of voxels within the manually traced mask for NM-1.5 mm sequence and SPM12 spatial normalization software, which is the worst performing method as per A. Yellow denotes excellent reliability (ICC over 0.75), orange denotes good reliability (ICC between 0.75 and 0.6), red denotes fair reliability (ICC between 0.6 and 0.4), and burgundy denotes poor reliability (ICC under 0.4).

### 3.5. Effect of spatial smoothing

A plot showing the relationship of ICC_ASV_ and CNR_V_ of voxels within the manually traced mask at 4 levels of spatial smoothing is shown in Fig. 8. Greater amounts of spatial smoothing lead to significantly lower CNR_V_ and significantly higher ICC_ASV_ (Wilcoxon signed rank test, *P* < 0.001 for all after correction for multiple comparisons). Relative to the one-lower degree of spatial smoothing (e.g., 2 mm vs 1 mm), spatial smoothing with 1 mm FWHM achieved the greatest increase in ICC_ASV_ and lowest decrease in CNR_V_, 0.03 and −0.09, respectively. Although there was still a significant difference in ICC_ASV_ and CNR_V_ between spatial smoothing with FWHM of 0 mm and 1 mm, the minimal difference in CNR_V_ and overall improvement in the robustness of voxelwise-analysis and spatial normalization in particular, support the use of spatial smoothing with 1 mm FWHM.

**Fig. 8:**
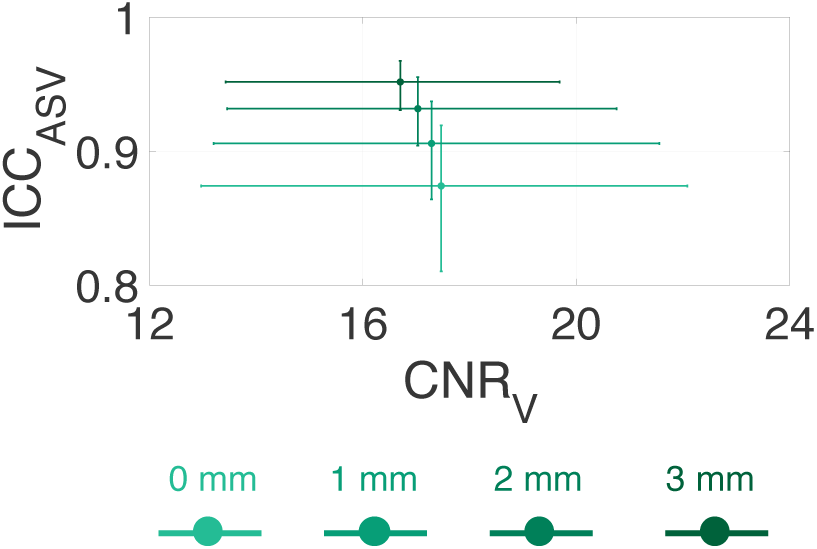
The effect of spatial smoothing on ICC_ASV_ and CNR_V_ of voxels within the manually traced mask of the SN/VTA-complex (Fig. 3B) for different degrees of spatial smoothing. Data points denote the median and error bars show the 25^th^ and 75^th^ percentiles.

### 3.6. Analyses using probabilistic atlas of dopaminergic nuclei

Using a recent high-resolution probabilistic atlas that identified the SN/VTA-complex nuclei (Pauli et al., 2018), we evaluated the feasibility of obtaining reliable measures of NM-MRI signal in these nuclei. Such measures would be valuable for basic and clinical neuroscience, particularly for the VTA, given its importance for reward learning (Montague et al., 1996; Schultz, 1998; Ungless et al., 2004) and affective processing (Depue and Collins, 1999; Fields et al., 2007). Plots of the ICC_ROI_ and CNR_ROI_ within the probabilistic masks as a function of acquisition time for the NM-1.5 mm sequence and ANTs spatial normalization software and various probability cutoffs are shown in Fig. 9. In general, excellent test-retest reliability was achieved for all nuclei and all probability cutoff values within 6 minutes of acquisition time. Similar to CNR in the manually traced mask, the CNR_ROI_ was not affected by acquisition time. The highest CNR was consistently observed in the SNr and SNc, then the PBP, and the lowest CNR in the VTA.

**Fig. 9:**
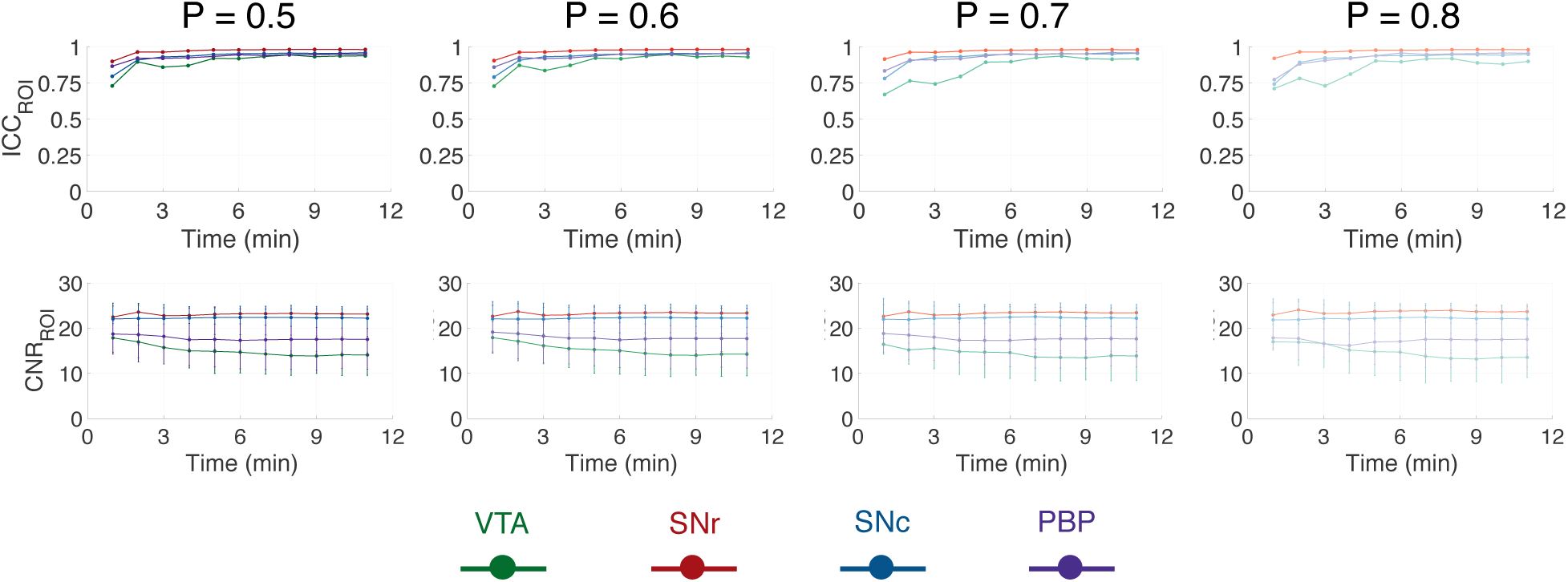
ICC_ROI_ (top row) and CNR_ROI_ (bottom row) within the probabilistic masks of the SN/VTA-complex nuclei (Fig. 3D) with different probability cutoffs (0.5, 0.6, 0.7, and 0.8) as a function of acquisition time. Data points denote the median and error bars indicate the 25^th^ and 75^th^ percentiles.

Having established the ability to reliably measure NM-MRI within SN/VTA-complex nuclei, we then investigated how distinct the NM-MRI signal within each nuclei is. The SN/VTA-complex nuclei are believed to have distinct anatomical projections and functional roles, so independently measuring signals from these nuclei would allow for investigation into these distinct anatomical circuits and functions. Although the nuclei are anatomically distinct, the potential for cross-contamination of the NM-MRI signal exists due to partial volume effects and spatial blurring due to MRI acquisition and the spatial normalization procedure. The independence of CNR_ROI_ values measured within the individual SN/VTA-complex nuclei was assessed by nonparametric Spearman’s correlation (Fig. 10). Overall, CNR was highly correlated across the four nuclei, particularly for ROI definitions based on a probability cutoff of P = 0.5.

**Fig. 10:**
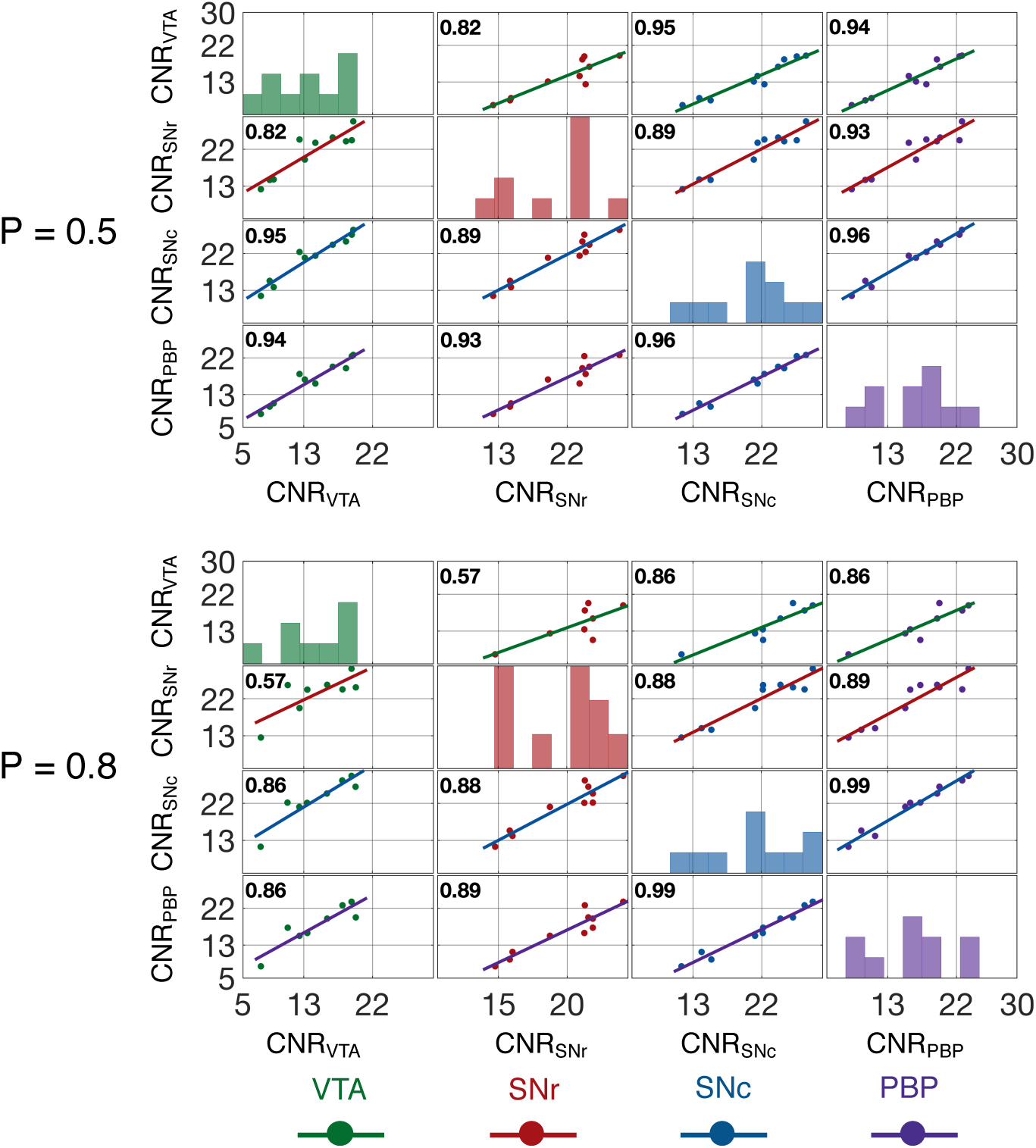
Correlations and histograms of the CNR_ROI_ values within the 4 SN/VTA-complex nuclei (Fig. 3D) for the lowest (P = 0.5) and highest (P = 0.8) probability cutoffs. The value within each correlation plot is Spearman’s rho.

## 4. Discussion

We present here a detailed description of a volume placement protocol for NM-MRI and use a test-retest study design to quantitatively derive recommendations for NM-MRI sequence parameters and preprocessing methods to achieve reproducible NM-MRI for ROI and voxelwise analyses. Additionally, by using a high-resolution probabilistic atlas, we were able to determine the reproducibility of NM-MRI measurements in specific nuclei within the SN/VTA-complex. Overall, excellent reproducibility was observed in all ROIs investigated and for voxels within the ROIs. Based on our results, we recommend acquiring at least 6 minutes of data for voxelwise-analysis or dopaminergic-nuclei-ROI-analysis and at least 3 minutes of data for standard ROI-analysis. We also recommend acquiring NM-MRI data with 1.5 mm slice-thickness, using ANTs for spatial normalization, and performing spatial smoothing with a 1 mm FWHM 3D Gaussian kernel for voxelwise-analysis and no spatial smoothing for ROI-analysis (especially for analysis of nuclei).

The goal of this paper was to provide detailed guidelines covering acquisition through analysis and recommendations for performing NM-MRI experiments with high quality and reproducibility. The main metric used to determine the recommendations was ICC. The high ICC values observed in our test-retest study suggest that NM-MRI using 2D GRE-MT sequences achieve excellent reproducibility across several acquisition and preprocessing combinations. This is in agreement with a previous report that observed an ICC_ROI_ value of 0.81 for SN in 11 healthy subjects (Langley et al., 2017). Surprisingly the present study observed higher ICC_ROI_ values (∼0.92) even though our study used a template defined SN mask and the two MRI scans were separated by 13 ± 13 days instead of using a subject-specific semi-automated thresholding method for SN mask generation (Chen et al., 2014) and having test-retest scans within a single session in one day (in which subjects were removed from the scanner after the first session, repositioned on the table, and scanned again). The improved ICC_ROI_ observed in our study could be explained by the rigor of the NM-MRI volume placement protocol that makes use of easy to identify anatomical landmarks to improve the reproducibility of volume placement across sessions. The study by Langley et al. focused on ROI measures and did not measure voxelwise ICC, so our study also extends this previous work in suggesting that voxelwise CNR measures can be obtained reliably. Another recent study measured ICC_ASV_ in 8 healthy and 8 patients with schizophrenia, also with both MRI sessions on the same day (∼1 hour apart) (Cassidy et al., 2019). That study observed a median ICC_ASV_ value of 0.64 and an ICC_ROI_ value of 0.96. The higher ICC_ASV_ observed in the present study could be due to the inclusion of only healthy subjects as well as the detailed volume placement protocol.

In addition to ICC values, we also used the strength of the NM-MRI signal (CNR) and the range of CNR values. Because correlation-based approaches are common for voxelwise-analysis, a greater range in CNR values within the SN/VTA-complex will provide greater statistical power. Another important factor in our analysis was the relationship between CNR and ICC. To make sure that voxelwise-analysis effects are not driven solely by high (or low) CNR voxels due to lower measurement noise in those voxels, it is important to have homogeneously high ICC values independent of CNR. Our recommended ANTs-based method applied to NM-MRI data with 1.5 mm slice-thickness accomplishes this independence.

This study is the first NM-MRI study to measure CNR in nuclei within the SN/VTA-complex. This was facilitated by making use of a publicly available high-resolution probabilistic atlas (Pauli et al., 2018). We demonstrated that the NM-MRI signal within the nuclei is highly reproducible with ∼ 6 minutes of data. Overall, we observed the highest CNR in the SNc and SNr, followed by the PBP, then the VTA. This is consistent with reports of higher degree of NM pigmentation in the SN than the VTA (Hirsch et al., 1988; Liang et al., 2004). However, we observed that the NM-MRI signal was highly correlated across nuclei. This finding may suggest that NM-MRI may only provide a measure of the general function of the dopamine system and is not specific to nuclei with distinct anatomy and function. While this may be true, our study included a limited number of subjects. Additionally, it is possible that the different functional domains of the dopamine system are highly correlated in healthy subjects and small errors in the realignment and spatial normalization processes could cause signal from different nuclei becoming mixed. These concerns could be partially mitigated through the use of a voxelwise-analysis (Cassidy et al., 2019). A future study should be conducted to investigate variability in specific functional domains of the dopamine system to determine if NM-MRI is capable of independently assessing dopamine functions more closely related to each of the nuclei.

This study tested 2 NM-MRI sequences with 3 mm slice-thickness: NM-3 mm and NM-3 mm Standard. The main difference between these two NM-MRI sequences was the use of in-plane acceleration, the number of slices, the TE, and the TR. These parameters were changed relative to the literature standard protocol (i.e. NM-3 mm Standard) to accommodate the increased number of slices required for similar coverage in the higher resolution sequences (i.e. NM-1.5 mm and NM-2 mm). Although the higher resolution sequences did not seem to be affected, increased noise due to in-plane acceleration could have caused the lower reproducibility observed for the NM-3 mm sequence compared to the NM-3 mm Standard sequence (Robson et al., 2008). An alternative explanation is that the reduced number of slices results in reduced performance of the realignment and coregistration steps (resulting from less anatomical information for the algorithms to work with) leading to reduced reproducibility. All images were manually inspected at each step and no obvious errors occurred but small-scale deviations in the preprocessing could impact the reproducibility. Future studies are needed to better understand the effect of TE and TR on the reproducibility of the NM-MRI signal.

This work provides a foundation for several applications in both basic and clinical neuroscience. The high reliability of NM-MRI observed is this study provides strong evidence for NM-MRI as a reliable noninvasive tool to investigate the role of the dopaminergic system in vivo. This includes the study of reward related behavior (Schultz, 2007) and investigation of dopaminergic abnormalities in addiction (Kelley and Berridge, 2002; Koob et al., 1998), Parkinson’s disease (Fahn and Sulzer, 2004; Sulzer et al., 2018; Sulzer and Surmeier, 2013), depression (Dunlop and Nemeroff, 2007; Nestler and Carlezon Jr, 2006), and schizophrenia (Abi-Dargham et al., 2000; Davis et al., 1991; Laruelle et al., 1996; Weinstein et al., 2017). Our findings also support previous research using NM-MRI in Parkinson’s disease (Aiba et al.; Cassidy et al., 2019; Hatano et al., 2017; Huddleston et al., 2017; Isaias et al., 2016; Kitao et al., 2013; Matsuura et al., 2013; Ohtsuka et al., 2014; Ohtsuka et al., 2013; Sasaki et al., 2006; Wang et al., 2019; Wang et al., 2018; Xing et al., 2018) and schizophrenia (Cassidy et al., 2019; Shibata et al., 2008; Watanabe et al., 2014; Yamashita et al., 2016), and suggest a continuation of this line of work. The ability to acquire highly reproducible NM-MRI measurements for ROI-analysis with a 3-minute acquisition opens the possibility of applications to clinical populations that may not have been previously possible due to prohibitively long acquisition times. While the mechanism of NM-MRI contrast are not fully understood and its specificity in measuring NM concentration or dopamine neuron loss requires further investigation, our results suggest that this technique may provide a reliable tool for the development of neuropsychiatric biomarkers.

Limitations of the present study include not conducting an exhaustive search of MRI acquisition parameters (e.g. TR, TE) or MT parameters (e.g. flip angle, offset). Such a search would have made the present study untenable, so we focused on what we perceived as the most critical acquisition parameter: slice-thickness. Additionally, as the contrast mechanisms of NM-MRI signal are better understood (Trujillo et al., 2017), numerical simulations could be performed to optimize several acquisition parameters including TE, TR, and MT parameters. Another limitation was the inclusion of only relatively young and healthy subjects. As such, the ICC values in patients (e.g. psychosis, Parkinson’s) could be lower due to reduced compliance in the MRI, however increased range of CNR values in patients could result in higher ICC values.

## 5. Conclusions

We have empirically determined that highly reproducible NM-MRI can be achieved with at least 6 minutes of NM-MRI data for voxelwise-analysis or nucleus-specific analysis, and at least 3 minutes of NM-MRI data for ROI-analysis. Additionally, we recommend using 1.5 mm slice-thickness, ANTs spatial normalization software, and spatial smoothing with 1 mm FWHM 3D Gaussian kernel. This work provides a foundation for the optimization and standardization of NM-MRI, a promising MRI approach with growing applications throughout basic and clinical neuroscience.

## Acknowledgments

The authors would like to thank Drs. Xiaoping Hu and Jason Langley for consultation with implementing the partial k-space MT sequence. Pilot funding for MRI scanner time was provided by the SCAN Center at Stony Brook University.

## Data and code availability

The code and the data used to produce the results reported in the manuscript are available from the corresponding author upon reasonable request.

